# Glycosomal Aquaglyceroporin 1 Dual Role in Iron Homeostasis and Antimony Susceptibility in *Leishmania amazonensis*

**DOI:** 10.1101/2025.08.14.670269

**Authors:** Romario Lopes Boy, Ricardo Andrade Zampieri, Juliana Ide Aoki, Adriano Cappellazzo Coelho, Lucile Maria Floeter-Winter, Maria Fernanda Laranjeira-Silva

## Abstract

*Leishmania* parasites cause a spectrum of diseases known as leishmaniases and must acquire nutrients like iron while surviving host defenses. Aquaglyceroporin 1 (AQP1) is a membrane channel that, in *L. major*, localizes to the flagellum and mediates antimony uptake and cell-volume regulation. Here, we show that in *L. amazonensis* AQP1 is instead targeted to glycosomes and that its expression is modulated by iron availability. A CRISPR-Cas9–mediated knockout of AQP1 in *L. amazonensis* revealed its multifunctional importance. AQP1-null promastigotes displayed a significant growth defect, particularly under iron-depleted conditions, and were impaired in regulating cell volume under osmotic stress. The mutant parasites contained approximately 50% less intracellular iron than wild-type cells and showed an increase in total superoxide dismutase activity, underscoring a role for AQP1 in iron homeostasis and oxidative stress management. AQP1 deletion also markedly reduced virulence in murine macrophages and in infected mice. Strikingly, loss of AQP1 increased resistance to trivalent antimony (Sb^III^), a first-line antileishmanial drug. AQP1-knockout promastigotes showed a 70% increase in Sb^III^ EC_50_ and accumulated more Sb intracellularly than wild-type, suggesting an altered antimony handling. Altogether, *L. amazonensis* AQP1 is a glycosomal protein that links iron metabolism, osmoregulation, and antimony susceptibility. Its glycosomal targeting and multifaceted roles differ from those of AQP1 orthologs in other *Leishmania* species. These findings suggest the existence of additional antimony uptake mechanisms beyond AQP1, with implications for understanding drug resistance.

**Author Summary:** Leishmaniases are neglected tropical diseases caused by parasites that survive and multiply inside vertebrates’ cells. These parasites rely on hosts’ nutrients like iron and must resist both host defenses and treatment with toxic drugs such as antimony. We studied a protein called Aquaglyceroporin 1 (AQP1) in *Leishmania amazonensis*, a species that causes skin lesions in South America. Unlike related species, where AQP1 is found on the parasite’s surface, we discovered that in *L. amazonensis* AQP1 is located in an internal organelle called glycosome. By deleting this protein from the parasite, we found that it plays a crucial role in iron balance, sensitivity to antimony drugs, and the parasite’s ability to cause disease. Unexpectedly, parasites without AQP1 were more resistant to antimony but still accumulated high levels of the drug, suggesting that *Leishmania* has other ways of taking up antimony. Our findings challenge the assumption that all *Leishmania* species use the same strategies to survive, and highlight the need to understand species-specific differences when designing treatments or analyzing parasite biology.

## Introduction

Leishmaniases are a group of neglected tropical diseases caused by kinetoplastid parasites of the genus *Leishmania*. They affect millions of people worldwide and remain a major public health problem, especially in tropical and subtropical regions where poverty and limited access to healthcare prevail. Clinical manifestations range from self-limiting cutaneous lesions to destructive mucocutaneous forms and fatal visceral disease if left untreated. Current estimates indicate approximately 12 million people are infected across 98 countries in Africa, Asia, the Americas, and Europe, with nearly one million new cases reported annually [1]. Additionally, around one billion people living in endemic areas are at risk of infection. This situation is further aggravated by deforestation and human expansion into forested areas, which facilitate sand fly vectors adapting to domestic environments and contribute the urbanization of the disease [2].

*Leishmania* parasites have a heteroxenous life cycle, alternating between phlebotomine sand fly vectors and vertebrate hosts [3]. Within vertebrate hosts, the parasite proliferates as amastigotes inside macrophages and other cells. The survival of *Leishmania* in these cells depends on its ability to evade host immune defenses, especially within macrophages phagolysosomes where amastigotes confront adverse conditions such as low pH, oxidative stress, and restriction of essential nutrients like iron [4–6].

Iron is essential for *Leishmania*, supporting processes such as mitochondrial respiration and defense against reactive oxygen species. The parasite acquires iron as ferrous iron (Fe^+2^) and as heme [5]. Ferrous iron import is mediated by the *Leishmania* Iron Transporter 1 (LIT1), whereas heme uptake is mediated by heme transporters like *Leishmania* Heme Response-1 (LHR1) [7]. Maintaining iron homeostasis is crucial since excess iron can catalyze the formation of harmful reactive species, leading to cytotoxicity [8]. To prevent toxic iron accumulation, *Leishmania* rely on the iron exporter *Leishmania* Iron Regulator 1 (LIR1), a plasma membrane protein shown to mediate iron efflux [9].

Trypanosomatids also possess unique peroxisome-related organelles called glycosomes that compartmentalize glycolysis and various other metabolic pathways, including purine salvage, pyrimidine biosynthesis, the pentose phosphate pathway, gluconeogenesis, fatty acid β-oxidation, ether-lipid biosynthesis, and oxidant defense [10, 11]. Some of those pathways require iron-dependent enzymes; however, the proteins responsible for iron trafficking in glycosomes remain unknown. The previous transcriptomic analysis of *L. amazonensis* under iron-deprived conditions identified several differential expressed genes encoding predicted transmembrane proteins with canonical peroxisomal targeting signals (PTS) [12]. One of these is the gene encoding a putative aquaglyceroporin-like protein. Aquaglyceroporins are members of the major intrinsic protein (MIP) superfamily, which form channels that facilitate the transport of water, glycerol and certain metalloids across biological membranes [13]. In *L. major*, an aquaglyceroporin 1 (AQP1) localized to the flagellum has been implicated in the uptake of trivalent antimony (Sb^III^) and in antimonial drug resistance [14, 15].

Here, we report that *L. amazonensis* AQP1 contains a C-terminal peroxisomal targeting sequence type 1 (PTS1) and is targeted to glycosomes rather than the flagellum. Using a CRISPR/Cas9 knockout approach combined with phenotypic analyses, we demonstrate that AQP1 contributes to the regulation of intracellular iron content, osmotic balance, parasite growth, infectivity, and susceptibility to antimonials. These findings position AQP1 as a dual regulator of iron trafficking and metalloid transport, revealing a previously unrecognized link between glycosomal physiology, iron homeostasis, and drug response in *Leishmania*.

## Results

### *L. amazonensis* AQP1 is a glycosomal protein and its expression is modulated during the parasite life cycle

Re-examination of a whole-genome transcriptome profile of *L. amazonensis* under iron deprivation indicated that gene LmxM.30.0020 is iron-responsive (fold-change = -1.2672; p = 0.043) [12]. This gene is annotated as Aquaglyceroporin 1 in *L. mexicana* (PMID:16135234) and is conserved across the genus *Leishmania*, with potential orthologs also found in *Crithidia fasciculata* (TriTrypDB CFAC1_270006300.1), *Endotrypanum monterogeii* (TriTrypDB EMOLV88_310005100.1), *Leptomonas pyrrhocoris* (TriTrypDB rna_LpyrH10_32_1110), *Leptomonas seymouri* (TriTrypDB rna_Lsey_0007_0780-1), and *Porcisia hertigi* (TriTrypDB JKF63_03741_t1) [16]. The ortholog in *L. amazonensis* is annotated as a putative major intrinsic protein (MIP) of 314 amino acids with six predicted transmembrane domains (molecular mass ∼34.778 kDa), which we hereafter refer to as *L. amazonensis* Aquaglyceroporin 1 (AQP1).

Proteins of the MIP superfamily form membrane channels that transport water, small neutral solutes, and in some cases metalloids. In *L. major*, AQP1 has been associated to drug resistance by mediating the efflux of trivalent antimony (Sb^III^) [17]. Characterization of *L. major* AQP1 showed that the protein localizes to the promastigote flagellum and, in amastigotes, to both the flagellar pocket membrane and contractile vacuole. This previous study also demonstrated AQP1’s role in solute transport, volume regulation, and osmotaxis [14]. To extend these findings, we generated a three-dimensional model of *L. amazonensis* AQP1 with AlphaFold 3 [18]. The fully automated modeling, including homology search and multiple sequence alignment, returned per-residue confidence scores (pLDDT), with values below 70 interpreted cautiously. The predicted fold and subunit interface showed high confidence, with pTM and ipTM scores of 0.81, indicating a robust overall structure and inter-chain interaction (Fig. 1a).

**Figure 1.**
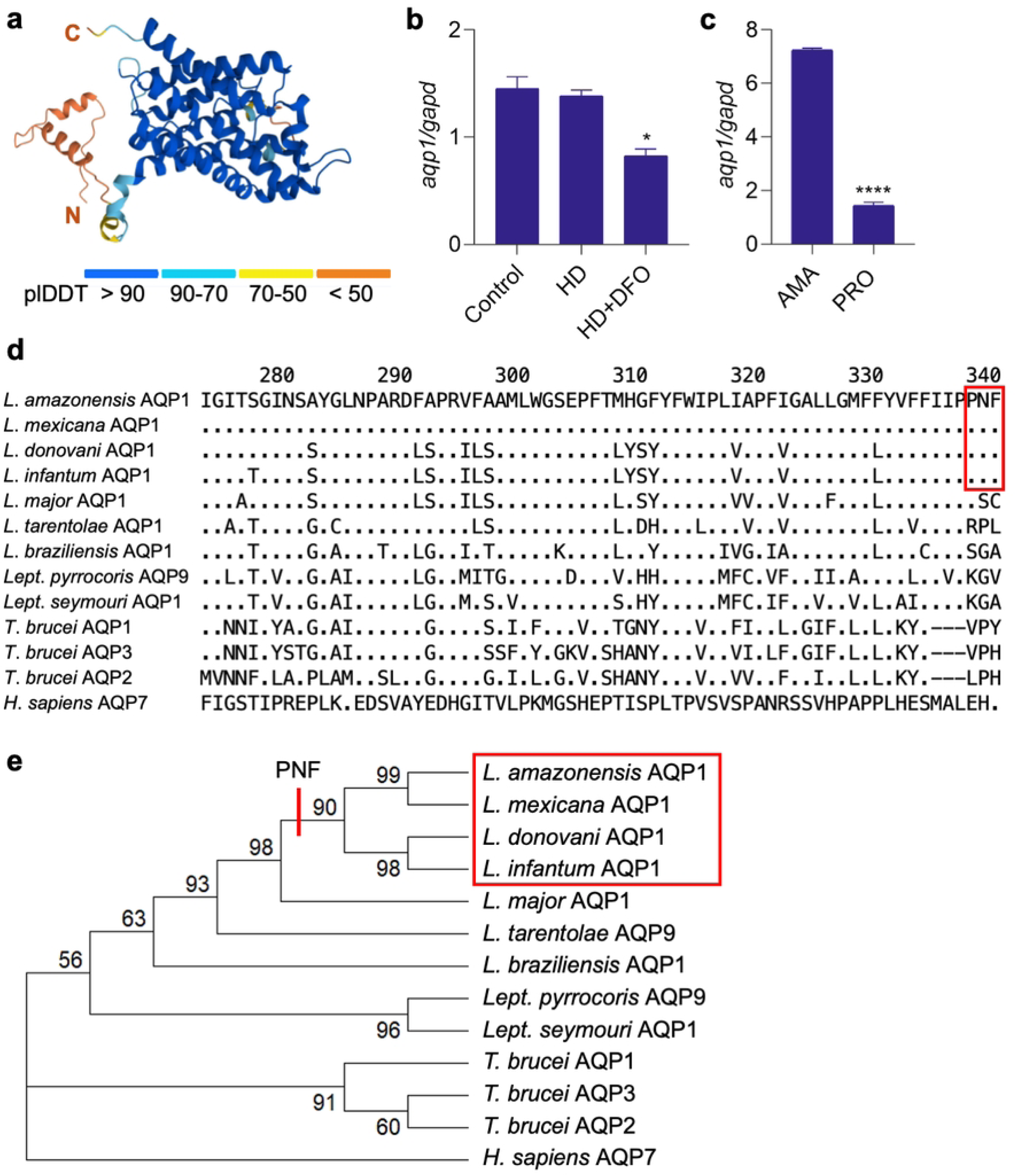
*L. amazonensis* AQP1 is encoded by an iron-responsive gene containing a glycosomal targeting signal. (a) AlphaFold model of *L. amazonensis* AQP1 showing six predicted transmembrane helices; colors reflect per-residue confidence (pLDDT score). N, amino-terminus; C, carboxi-terminus. (b) *aqp1* transcript levels in promastigotes grown in complete medium (Control) or under iron-restricted conditions: heme-depleted medium (HD) alone or with 50 µM deferoxamine (HD + DFO). (c) *aqp1* transcript levels in axenic amastigotes and promastigotes. Promastigote transcript levels were determined daily from day 2 to day 7 of culture (bar shows the average across these time points). Transcript levels were quantified by RT-qPCR and normalized to the *glyceraldehyde-3-phosphate dehydrogenase* (*gapd*) reference gene. Bars represent mean ± SEM of two independent experiments. *p <0.05; ****p <0.0001 (one-way ANOVA with Dunnett’s post-test). (d) Multiple alignment of the C-terminal 68 amino acids of trypanosomatid aquaglyceroporins. The Pro-Asn-Phe (PNF) peroxisomal-targeting motif is boxed in red. Dots (.) denote residues identical to the reference (top) sequence; dashes (–) indicate gaps. (e) Maximum-likelihood phylogeny of the same proteins. Bootstrap values (1,000 replicates) are shown at each node. Branches containing the PNF/PTS1 motif are highlighted with a red bar. *Homo sapiens* AQP7 was used as an outgroup. Protein IDs: *L. amazonensis* LAMAPH8_000653100; *L. mexicana* LmxM.30.0020; *L. donovani* LdBPK_310030; *L. infantum* LINF_310005100; *L. major* LmjF.31.0020; *L. tarentolae* LtaP31.0020; *L. braziliensis* LbrM.00.0079; *Trypanosoma brucei* Tb927.6.1520, Tb927.10.14160, Tb927.10.14170; *Leptomonas pyrrhocoris* LpyrH10_32_1110; *Lept. seymouri* Lsey_0007_0780; *H. sapiens* NP_001161.1.

Quantitative RT-PCR analysis confirmed that *aqp1* transcript levels are downregulated when promastigotes are cultured under iron-poor conditions. Specifically, after 18 hours in heme-depleted medium supplemented with the iron chelator deferoxamine (DFO), *aqp1* mRNA levels were significantly lower than those in iron-replete control cells (Fig. 1b). Besides, we observed a 5-fold increase in *aqp1* expression in axenic amastigotes compared to the average transcript levels in promastigotes (Fig. 1c).

Sequence analysis of *L. amazonensis* AQP1 revealed the presence of a canonical PTS1 motif (Pro-Asn-Phe, “PNF”) at the C-terminus of the protein (Fig. 1d). This tripeptide is a well-characterized glycosomal targeting signal [11, 19]. Notably, alignment of AQP1 orthologues from various trypanosomatids showed that the PTS1 motif is present only in *L. amazonensis*, *L. mexicana*, *L. donovani*, and *L. infantum*. This motif is absent from AQP1 in other *Leishmania* species (Fig. 1d), including *L. major* AQP1, whose AQP1 localizes to the flagellum, consistent with the lack of PTS1 (Figarella et al., 2007). Pairwise alignment indicated ∼74.8 % amino acid identity between the *L. amazonensis* AQP1 and *L. major* AQP1 (Supplementary Fig. 1). Moreover, a maximum-parsimony phylogenetic tree revealed that the species harboring the PTS1 motif in AQP1 form a monophyletic clade, comprising *L. amazonensis*, *L. mexicana*, *L. donovani* and *L. infantum*, distinct from other *Leishmania* species (Fig. 1e).

To verify the PTS1-based prediction, we generated a transgenic *L. amazonensis* line expressing AQP1 tagged with EGFP at the N-terminus. Confocal immunofluorescence revealed EGFP–AQP1 colocalization arginase, a glycosomal marker [20] (Fig. 2), confirming glycosomal targeting. In contrast, previous studies reported different AQP1 localizations in other species: cell surface/flagellum in *L. major* [14, 21], and an undefined cytosolic signal in *L. donovani* [22]. Together, these data demonstrate that AQP1 is glycosomal in *L. amazonensis*. Moreover, the downregulation of *aqp1* under iron deprivation and its upregulation in infective amastigotes support a role in intracellular iron trafficking, linking glycosomal physiology to iron homeostasis.

**Figure 2.**
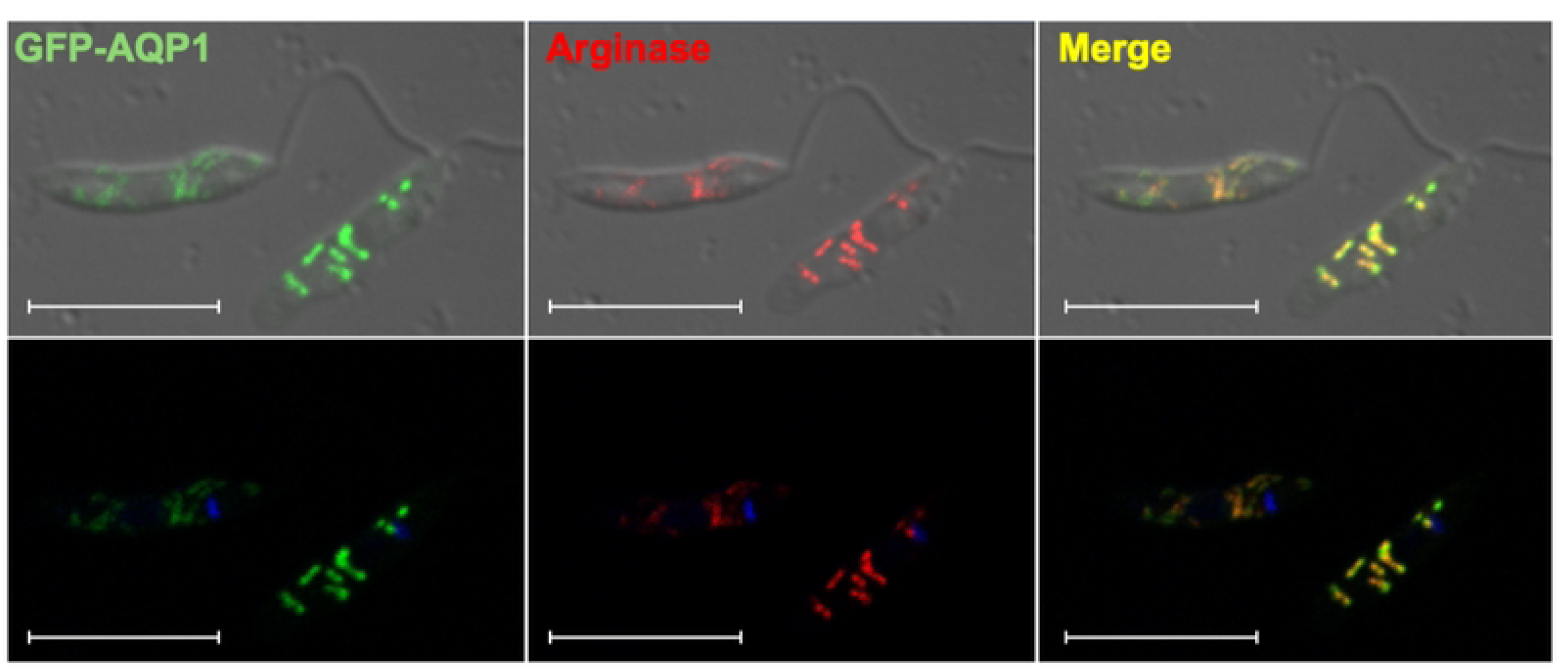
*L. amazonensis* AQP1 localizes to glycosome in promastigotes. Confocal immunofluorescence images of *L. amazonensis* promastigotes expressing EGFP-AQP1. Green: EGFP-AQP1 fusion; Red: glycosomal marker (anti-arginase); Yellow: merged image; Blue: DNA (DAPI). Scale bar = 10 µm.

### Loss of AQP1 impairs promastigote growth under heme-depleted conditions

To investigate AQP1’s role in *L. amazonensis*, we generated homozygous *aqp1* knockout mutant lines (*aqp1-/-*) using the LeishGEdit CRISPR/Cas9 system [23, 24]. First, an *L. amazonensis* PH8 line ectopically expressing Cas9 nuclease and T7 RNA polymerase (C9/T7) was obtained and validated. Western blotting confirmed Cas9 expression in this line, and the C9/T7 parasites showed no growth defects or changes in antimony susceptibility or virulence compared to wild-type controls (Supplementary Fig. 2). We then transfected C9/T7 promastigotes with a donor DNA construct containing a puromycin resistance gene (*pac*) flanked by 5′ and 3′ homology arms, along with two single-guide RNA templates targeting sequences immediately upstream and downstream of the *aqp1* coding region (Fig. 3a). Although *aqp1* resides on the tetrasomic chromosome 30 [25], a single round of CRISPR editing was sufficient to knock out all alleles. We obtained two independent knockout clones (KO1 and KO2) in which the *aqp1* open reading frame was replaced by the *pac* cassette, as confirmed by PCR (Fig. 3b). For genetic complementation, we introduced an episome containing *aqp1* ORF tagged with EGFP into the KO1 clone, generating add-back lines (AB1–AB4) that were also verified by PCR (Fig. 3b). The KO1 clone and AB1 clone were selected for further phenotypic analyses.

**Figure 3.**
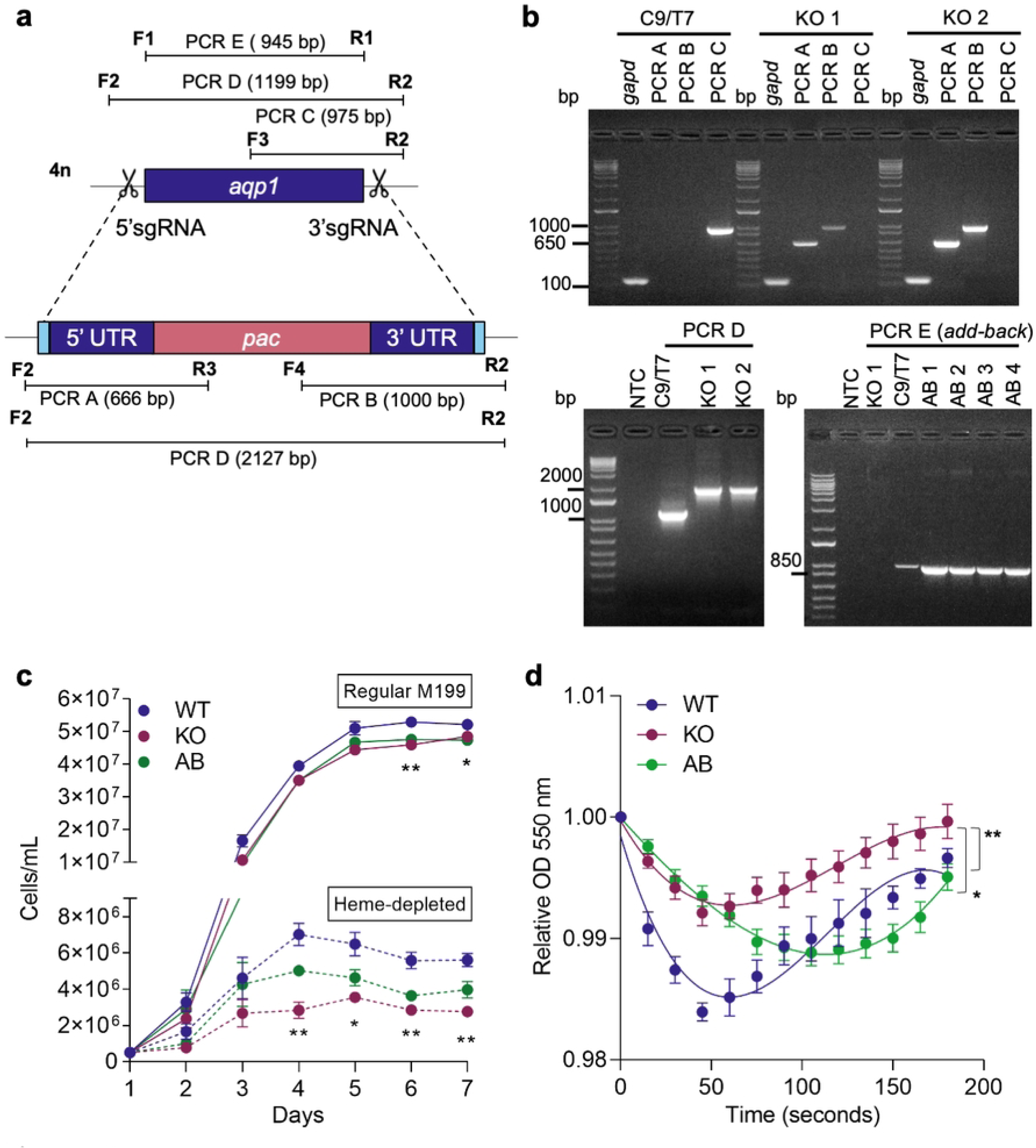
Impact of AQP1 loss on promastigote replication and osmoregulation. (a) Schematic of the CRISPR/Cas 9 strategy to delete *aqp1* in *L. amazonensis*. Two sgRNAs (scissors) directed Cas9 to cut at 5′ and 3′ sites flanking the *aqp1* ORF. A donor DNA containing a puromycin resistance marker (*pac*) with flanking 5′ and 3′ untranslated region (UTR) homology arms were delivered to replace the *aqp1* ORF via homology-directed repair. Black bars indicate primer binding sites (F1–F4 and R1–R3) used for diagnostic PCR, producing amplicons PCR A–PCR E of the indicated sizes in base pairs (bp). (b) PCR confirmation of *aqp1* deletion and complementation. Agarose gel images show diagnostic PCR products for the parental Cas9/T7 line (C9/T7), two *aqp1* knockout clones (KO1, KO2), and four add-back clones (AB1–AB4). Water was used as a no-template control (NTC). DNA ladder sizes in bp. *Gapd* = control PCR targeting the *glyceraldehyde-3-phosphate dehydrogenase* gene (indicating successful DNA amplification for each template). (c) Growth curves of WT, *aqp1* knockout (KO), and add-back (AB) promastigotes in regular medium (solid lines) vs. heme-depleted medium (dashed lines). Parasite density (cells/mL) is shown as mean ± SEM of three independent experiments. * p < 0.05; ** p < 0.005 (KO vs. WT). (d) Osmotic-stress responses of WT, KO, and AB promastigotes, measured as changes in optical density over 3 min following a hypo-osmotic shock. Data are mean ± SEM of three independent experiments. A non-linear sigmoidal fit (third-order polynomial) was applied, followed by one-way ANOVA and Tukey’s multiple comparisons test. **p = 0.0034 (WT vs. KO); *p = 0.0457 (KO vs. AB).

We next compared the *in vitro* growth of wild-type (WT), AQP1-knockout (KO), and add-back (AB) promastigotes under normal and iron-limited conditions. Loss of AQP1 caused a significant growth defect, most evident in heme-depleted medium (Fig. 3c). As cultures progressed from logarithmic to stationary phase, KO parasites consistently reached lower densities than WT and AB, with the disparity becoming even more pronounced under heme deprivation.

Because AQP1 has been implicated in osmoregulation in *L. major* [21], we assessed whether loss of AQP1 affected the ability of *L. amazonensis* promastigotes to cope with osmotic stress. Cell volume changes in response to a hypo-osmotic shock were measured for WT, KO, and AB lines (Fig. 3d). KO cells exhibited a markedly smaller volume increase upon osmotic challenge compared to WT, indicating an impaired capacity for osmotic regulation in the absence of AQP1. The AB line showed partial restoration of the WT phenotype, which is a common outcome for episomal add-backs in *Leishmania* [9, 26, 27]. These results reveal a previously unrecognized role for AQP1 in promoting promastigote replication, especially under iron-poor conditions, and reinforce its involvement in maintaining osmotic balance.

### Glycosomal AQP1 impacts susceptibility to antimony and regulates intracellular iron levels

*Leishmania* susceptibility to antimonials has been linked to AQP1 expression, which would be involved in antimony uptake [28–30]. We therefore tested whether AQP1 deletion affects Sb^III^ susceptibility in *L. amazonensis*. MTT assays revealed that KO promastigotes were significantly more resistant to Sb^III^ than WT (Fig. 4a). The EC_50_ for Sb^III^ was 315 µM in the KO, 187 µM in the WT, and 121 µM in the AQP1 add-back line. Notably, the *L. amazonensis* WT strain was ∼7-fold more resistant than a *L. major* WT strain (EC_50_ 27 µM), consistent with species-specific differences in drug susceptibility [15]. We next measured antimony uptake by inductively coupled plasma mass spectrometry (ICP-MS) after exposing promastigotes to Sb^III^ for 15 minutes (Fig. 4b). Surprisingly, despite its higher resistance, the AQP1 KO line accumulated more Sb than WT parasites. This finding suggests that the presence of AQP1 may actually limit net Sb uptake, promote efflux reducing the intracellular accumulation of antimony, or detoxify Sb in *L. amazonensis*.

**Figure 4.**
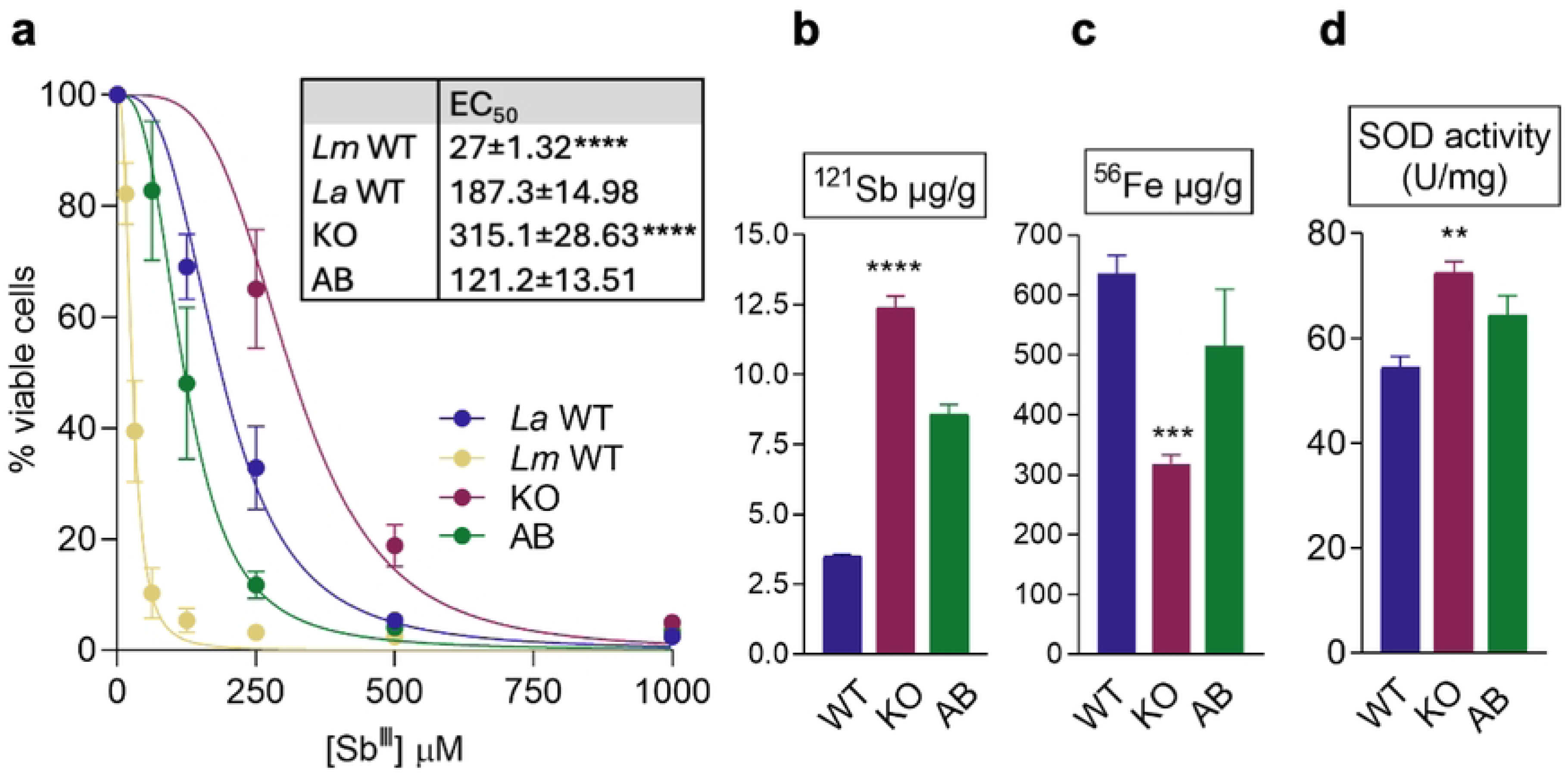
AQP1 mediates antimony susceptibility, intracellular iron levels, and SOD activity. (a) MTT viability assays for promastigotes of *L. major* wild type (*Lm* WT), *L. amazonensis* wild type (*La* WT), *aqp1* knockout (KO), and add-back (AB) lines exposed to increasing Sb^III^ concentrations (0–1000 µM). Percent survival is plotted for each strain. EC_50_ values are listed in the table (inset). Data are mean ± SEM from 3 independent experiments (each in triplicate). ****p<0.0001. (b) Total antimony content in WT, KO, and AB promastigotes after 15 min exposure to potassium antimonyl tartrate trihydrate (Sb^III^), measured by ICP-MS. Values are normalized to total cellular protein. Bars show mean ± SEM of 3 independent experiments (each in duplicate). **** p<0.0001. (c) Total iron content in WT, KO, and AB promastigotes at end-log growth phase, measured by ICP-MS. Values are normalized to total protein. Bars show mean ± SEM of 3 independent experiments (each in duplicate). ***p<0.001. (d) Total superoxide dismutase (SOD) activity in WT, KO, and AB promastigote lysates. The SOD activity is represented as units (U)/mg protein. Bars represent mean ± SEM of 3 independent experiments. ** p<0.01.

Given the iron-responsive expression of *aqp1* and the growth defect of the KO under iron-poor conditions, we examined whether AQP1 influences iron homeostasis. We quantified total cellular iron in WT, KO, and AB promastigotes by ICP-MS (Fig. 4c). The AQP1 KO contained roughly 50% less iron than WT cells, whereas iron levels in the AB line were restored to near-WT values. These results indicate that AQP1 is important for maintaining normal iron content in *L. amazonensis*. Given that trypanosomatids SOD enzymes utilize iron as a cofactor (unlike most eukaryotic SODs, which use manganese) and that the essential SOD B isoform is glycosomal in *Leishmania*, we also examined SOD expression and activity in the AQP1 mutant lines. Immunoblotting detected no differences in SODA or SODB protein levels between WT, KO, and AB parasites, regardless of heme availability (Supplementary Fig. 3). However, total SOD enzymatic activity was significantly elevated in the AQP1 KO compared to WT (Fig. 4d), indicating a compensatory upregulation of SOD activity in response to AQP1 loss (potentially through post-translational mechanisms).

Together, these data highlight a novel interesting link between antimony resistance and iron metabolism in *L. amazonensis*. Glycosomal AQP1 appears to be a key factor connecting these pathways, influencing both metal ion homeostasis and the oxidative stress response.

### AQP1 is essential for intracellular proliferation and *in vivo* pathogenicity

Infection of murine bone marrow–derived macrophages demonstrated that AQP1 is critical for the intracellular replication of *L. amazonensis* amastigotes (Fig. 5a). At 4 hours post-infection, similar numbers of WT, KO, and AB parasites were observed inside macrophages, indicating that AQP1 deletion does not impair initial host cell entry. However, by 48 hours post-infection, the number of KO amastigotes per macrophage had significantly decreased, whereas WT and AB parasites continued to replicate, indicating that KO parasites fail to proliferate intracellularly.

**Figure 5.**
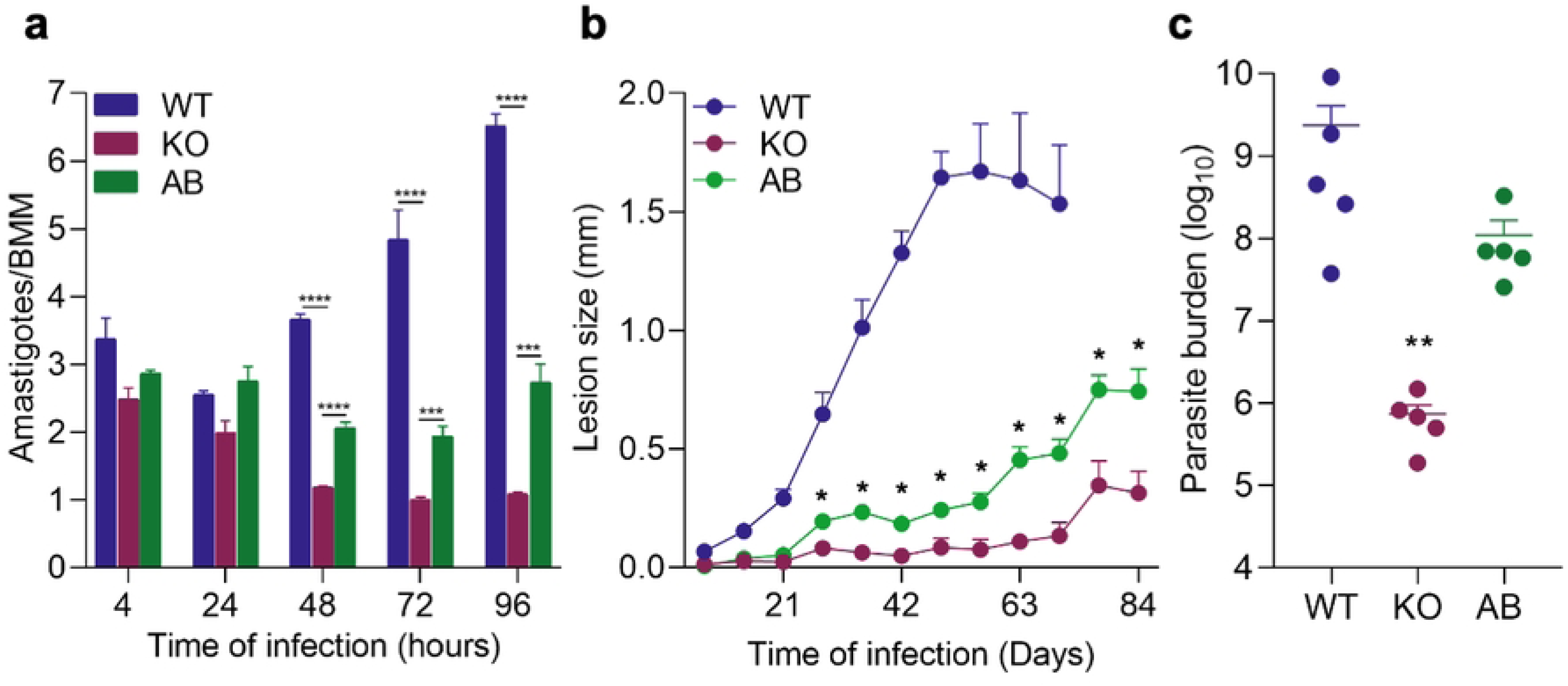
AQP1 deficiency reduces *L. amazonensis* infectivity in macrophage and mice. (a) Quantification of intracellular amastigotes in murine bone marrow–derived macrophages (BMM) infected with WT, KO, or AB metacyclic parasites (MOI = 5). Bars indicate the mean number of amastigotes per macrophage (± SEM) at 4, 24, 48, 72, and 96-hours post-infection from three independent experiments. *** p<0.001; ****p<0.0001. (b) Lesion development in C57BL/6 mice inoculated with 10^6^ metacyclic promastigotes of WT, KO, and AB lines. Points represent mean footpad thickness (± SEM, n = 5 mice per group) over time. * p ≤ 0.053 for AB vs. KO from days 24 to 84 post-infection (t-test). (c) Parasite load in footpad lesions at 10 weeks post-infection for WT (when lesions reached the maximum allowed) and 12 weeks for KO and AB (when lesions became apparent). Bars show mean ± SEM for 5 mice per group. ** p = 0.0079 (WT vs. KO; KO vs. AB).

We next examined the importance of AQP1 for *L. amazonensis* virulence *in vivo*. C57BL/6 mice were inoculated in the footpad with metacyclic promastigotes of WT, KO, or AB lines, and lesion development was monitored weekly. All mice infected with WT parasites developed progressive lesions by 3–4 weeks post-infection. In contrast, mice infected with AQP1 KO parasites showed only minimal pathology, with footpad swelling only becoming apparent around 11–12 weeks post-infection (Fig. 5b). Mice infected with AB parasites developed lesions of intermediate severity, with footpad thickness approximately two-fold greater than in the KO group. Parasite load measurements at the end of the experiment confirmed that KO-infected footpads harbored significantly fewer parasites than those infected with WT or AB lines (Fig. 5c).

These findings, together with the macrophage infection data, demonstrate that AQP1 is a pivotal virulence factor for *L. amazonensis*. Loss of AQP1 severely compromises the parasite’s ability to replicate inside host cells and cause disease in the mammalian host.

## Discussion

Our study reveals that aquaglyceroporin 1 (AQP1) in *L. amazonensis* is a glycosomal protein with multiple roles in parasite physiology. This discovery underscores how even well-conserved proteins can have divergent subcellular localizations and functions in different *Leishmania* species. In *L. major*, for instance, AQP1 localizes to the flagellum and flagellar pocket membranes, where it functions in osmoregulation and mediates antimony uptake [14, 31]; phosphorylation can further redistribute AQP1 over the entire parasite surface [21]. In contrast, we found that *L. amazonensis* AQP1 carries a C-terminal PTS1 motif (absent in *L. major* and several other species) and it is directed to glycosomes. To our knowledge, this is the first report of an aquaglyceroporin targeted to the glycosomal compartment in trypanosomatids [32]. Such species-specific differences are often overlooked. Large-scale projects, which map protein locations in a representative species, implicitly assume uniformity across the *Leishmania* genus, and sometimes even across the Trypanosomatidae family [33]. Our findings caution that generalizations from one species to all *Leishmania* can be misleading. Evolutionary adaptations, such as the PTS1 acquisition in AQP1 by species of the *L. mexicana* and *L. donovani* complexes, mean that protein localization and function may vary significantly, with important biological consequences. The discovery that *L. amazonensis* AQP1 resides in glycosomes establishes, for the first time, a direct connection between aquaglyceroporins and the peroxisome-like organelles of trypanosomatids, an association that until now had been only speculative [32]. Glycosomes compartmentalize key metabolic processes, including glycolysis, β-oxidation, and detoxification pathways, some of which require iron-dependent enzymes [34]. However, how iron and other solutes traffic into glycosomes remains largely unknown [5, 32]. We observed that *aqp1* expression is iron-responsive and is down-regulated under iron-depleted conditions. Furthermore, AQP1-null parasites retain only about half the intracellular iron of wild-type parasites.

These findings indicate that AQP1 helps maintain iron balance. One intriguing possibility is that AQP1 facilitates the movement of metabolites or small solutes crucial for storage or utilization within glycosomes. For instance, *Leishmania* encode a glycosomal superoxide dismutase (SODB) that uses iron as a cofactor [35, 36]; AQP1 might indirectly support such enzymes by maintaining osmotic and redox equilibrium inside glycosomes. In AQP1-null parasites, we observed no change in SOD protein levels but a significant increase in total SOD enzymatic activity, indicating a compensatory response to altered redox homeostasis. The impaired growth of AQP1-null mutant in iron-poor conditions further supports AQP1’s importance under iron-limited conditions, perhaps helping the parasite efficiently recycle or redistribute iron during scarcity. Overall, our results position glycosomal AQP1 as a factor in maintaining iron homeostasis, linking an aquaglyceroporin to metal metabolism in a way not previously appreciated.

Another major finding is that *L. amazonensis* AQP1 modulates sensitivity to antimonial drugs in an unexpected way. In other *Leishmania* species, AQP1 has been considered the principal route for uptake of trivalent antimony (Sb^III^), the active form of pentavalent antimonials [37]. Reduced AQP1 expression or loss-of-function mutations correlate with decreased Sb^III^ accumulation and drug resistance in *L. major* and in some clinical isolates. Indeed, wild-type *L. major* is far more Sb-sensitive than an AQP1-deficient line [17, 37]. Our *L. amazonensis* data, however, challenge this model. Deleting AQP1 in *L. amazonensis* did make parasites more Sb^III^-resistant (EC_50_ increased ∼1.7-fold), but paradoxically those AQP1-null parasites accumulated more Sb intracellularly than wild-type parasites. This indicates that Sb^III^ can still enter *L. amazonensis* cells by alternate routes when AQP1 is absent. One possibility is passive diffusion of Sb(OH)_3_ (the uncharged form of Sb^III^) across cell membranes. Normally, AQP1 greatly accelerates Sb(OH)_3_ uptake; in its absence, the compound may still slowly permeate the plasma membrane. Additionally, other transporters or channels might contribute to Sb uptake. Notably, in a highly Sb-resistant *L. guyanensis* strain lacking functional AQP1, Sb^III^ influx was not saturable, consistent with the presence of a low-affinity, non-specific uptake mechanism [38]. *L. amazonensis* AQP1-null mutants may exploit a similar secondary pathway.

Beyond uptake, the increased Sb content in AQP1-null parasites suggests that wild-type AQP1 might normally limit net Sb accumulation. Because *L. amazonensis* AQP1 is glycosomal, we speculate that it could facilitate sequestration or efflux of Sb. For example, AQP1 might transport Sb(OH)_3_ or a related metabolite into glycosomes, compartmentalizing Sb away from critical cytosolic targets or aiding its conversion into a less toxic form. In AQP1’s absence, more Sb remains in the cytosol, where it might be pumped into vesicles or otherwise neutralized [30]. Consistent with this idea, we found that AQP1-null parasites have a major perturbation in iron metabolism that likely affects their response to Sb. The knockout’s iron content was roughly half that of wild-type, which would reduce iron-driven Fenton reactions and the associated oxidative damage that Sb can exacerbate. In addition, AQP1-null mutants showed elevated SOD activity, which could improve clearance of Sb-induced reactive oxygen species. Antimonial drugs are known to trigger oxidative stress in *Leishmania*, directly or indirectly by depleting thiols [39], so parasites with bolstered antioxidant defenses and lower iron might better withstand this stress. Thus, changes in metal ion balance and redox environment in the AQP1 knockout provide a plausible explanation for how it tolerates higher Sb levels. Our findings reveal a previously unappreciated resistance mechanism: by altering iron homeostasis and the intracellular redox milieu, the parasite becomes less susceptible to Sb-induced damage.

This insight calls for a refinement of existing models of antimonial action and resistance, which have largely focused on drug import/export and thiol sequestration. Future studies could explore how manipulating parasite iron levels or oxidative stress pathways influences antimonial efficacy. It will also be important to identify the alternative Sb uptake route in AQP1-deficient *L. amazonensis*, whether through another aquaglyceroporin, a different type of channel, or simply passive diffusion, and to determine if similar pathways operate in other *Leishmania* species or in clinical scenarios.

Our data also show that AQP1 is required for optimal growth in host cells, underscoring its role as a virulence factor in *L. amazonensis*. AQP1 expression is significantly higher in the amastigote intracellular stage than in promastigotes, presumably to meet the parasite’s needs within the host. Inside phagolysosomes, amastigotes face nutritional immunity, such as iron deprivation, as well as osmotic and oxidative stresses [5]. AQP1 likely helps the parasite overcome these challenges, possibly by facilitating uptake of essential solutes, maintaining osmotic balance in the phagolysosomal compartment, or enabling efficient use of limited iron. Consistently, in the absence of AQP1, *L. amazonensis* amastigotes failed to proliferate in macrophages and produced only greatly attenuated lesions in mice.

Prior studies reported species-specific differences in AQP1 localization [14, 22]. In *L. major* amastigotes localize AQP1 to the flagellar pocket and contractile vacuole membranes. By contrast, we found that *L. amazonensis* AQP1 is glycosomal, so amastigotes of this species concentrate AQP1 internally in glycosomes rather than on the cell surface. Glycosomal AQP1 could thus be critical for metabolic flexibility in this species. Amastigotes glycosomes may need to rapidly adapt between nutrient-rich and nutrient-poor conditions, and AQP1 could facilitate fluxes of metabolites in and out of these organelles to sustain essential pathways. In the AQP1-null parasites, such flexibility appears lost, impairing the parasite’s ability to thrive within macrophages. This points to a metabolic vulnerability that could potentially be exploited for therapeutic intervention.

Taken together, our study demonstrates that *L. amazonensis* AQP1 serves as a dual regulator of iron homeostasis and antimony response, all while being tucked away in an organelle not previously associated with these processes. First, our findings reinforce that *Leishmania* species should not be treated as biologically interchangeable. Even highly conserved proteins can have subtle sequence differences, such as presence or absence of a PTS1 targeting motif, that redirect their localizations and function, significantly altering the parasite physiology. Researchers using reference strains and large datasets should integrate comparative analyses across species to avoid overlooking such unique adaptations. Examining multiple species will be key to understanding the full spectrum of strategies *Leishmania* employ for nutrient uptake and drug resistance. Second, our discovery opens new avenues to explore glycosomal functions and glycosome–cytosol communication. The glycosomal targeting of AQP1 appears to have evolved in specific lineages (including *L. donovani*, *L. infantum*, *L. mexicana*, *L. amazonensis*), suggesting that in these species glycosomes could play a more pronounced role in stress adaptation, surviving macrophage conditions or oxidative bursts for example. Finally, the link between metal ion homeostasis and drug susceptibility highlighted here suggests novel therapeutic angles. Manipulating parasite iron availability or redox balance might alter antimonial drug efficacy, pointing to new strategies to enhance treatment outcomes against leishmaniasis.

In conclusion, this work deepens our understanding of AQP1’s role in *Leishmania* biology. Rather than acting solely as a drug uptake channel, AQP1 in *L. amazonensis* emerges as a multifaceted protein at the crossroads of iron metabolism, osmotic stress adaptation, and drug resistance. These insights challenge assumptions drawn from other species and underscore the extent of evolutionary diversification within the *Leishmania* genus. Appreciating and dissecting such species-specific features will be crucial for developing more effective, targeted strategies to combat leishmaniasis across its many forms.

## Materials and Methods

### Animals

Female C57BL/6 mice (6–8 weeks old) (five animals per group) were obtained from the Animal Centre of the Medical School of the University of São Paulo and maintained at the Animal Centre of the Department of Physiology at the Institute of Bioscience of the University of São Paulo, receiving food and water *ad libitum*.

### Ethics statement

All animal experimental procedures were approved by the Animal Care and Use Committee at the Institute of Bioscience of the University of São Paulo (CEUA 342/2019) and were conducted in accordance with the recommendations and the policies for the Care and Use of Laboratory Animals of São Paulo State (Lei Estadual 11.977, de 25/08/2005) and the Brazilian government (Lei Federal 11.794, de 08/10/2008).

### *Leishmania* cultivation

Promastigote forms of the *L. amazonensis* IFLA/BR/67/PH8 and *L*. *major* MHOM/IL/81/Friedlin strains were cultured *in vitro* at 26 °C in M199: medium 199 (Gibco, Invitrogen) pH 7.2 supplemented with 10% heat inactivated fetal bovine serum (FBS), 40 mM HEPES, 0.1 mM adenine, 0.0001% biotin, 5 μg/ml hemin (25 mg/ml in 50% triethanolamine), 5 mM L-Glutamine and 5% penicillin-streptomycin. Heme-depleted medium was prepared similarly, omitting hemin addition and replacing regular FBS by heme-depleted FBS. Heme-depleted FBS was prepared as previously described [12] by treating heat inactivated FBS with 10 mM ascorbic acid for 16 h at room temperature, followed by verification of heme depletion by measuring the optical absorbance at 405 nm, 3 rounds of dialysis in cold phosphate-buffered saline (PBS) and filter-sterilization.

### *Leishmania* immunofluorescence microscopy

Promastigotes were fixed with 4% paraformaldehyde and attached to poly L-lysine coated slides. The fixed cells were quenched with 50 mM NH_4_Cl for 1 h and permeabilized with PBS containing 0.1% Triton X-100 for 15 min, prior to blocking with TBS Blocking Buffer (LI-COR Bioscience) for 1 h at room temperature. For arginase detection, a rabbit anti-arginase was used as primary antibody [20] (1:500 in blocking buffer) for 1 h, followed by an anti-rabbit AlexaFluor 546 (Life Technologies) (1:500) as secondary antibody. All samples were incubated with 1 μg/mL DAPI (4’,6-diamidino- 2-phenylindole) for nuclear staining. Slides were mounted with ProLong™ Gold Antifade Mountant (Invitrogen), and images were acquired using a confocal microscope (Zeiss LSM 780 NLO) at the Core Facility for Research Support (Centro de Facilidades para Pesquisa, CEFAP) from University of São Paulo.

### *aqp1* mRNA expression

Total RNA was obtained using PureLink RNA Mini Kit (Invitrogen) following manufacturers’ instructions. cDNA was synthesized using High-Capacity cDNA Reverse Transcription Kit (Applied Biosystems). For a final reaction volume of 20 μL, 2 μg of total RNA mixed with 0.5 μg oligo(dT)12-18, 0.8 μL of dNTPs 25 mM, 2 μL of Buffer 10X, and 50 U of Reverse Transcriptase (Applied Biosystems) was incubated at 25°C for 10 min, 37°C for 120 min, 85°C for 5 min, and storage at -20°C. A negative control containing all reaction components except the enzyme was included and analyzed by real-time PCR to exclude the possibility of DNA contamination in the RNA samples.

For real-time PCR, 1/20 of the reverse transcription product was used as a template. The reactions were performed in a StepOnePlus™ Real-Time PCR System (Applied Biosystems) with 200 nM of each gene-specific pair of primers and LuminoCt® SYBR® Green qPCR ReadyMix™ (Sigma Aldrich). The specific primers used were AQP1-RT-F and AQP1-RT-R for *L. amazonensis aqp1*, and GAPD-RT-F and GAPD-RT-R for *glyceraldehyde-3-phosphate dehydrogenase* (*gapd*) (S1 Table). The PCR reaction consisted of an initial denaturation step of 95°C for 20 s followed by 40 cycles of 3 s at 95°C and 30 s at 60°C. The target gene expression levels were quantified according to a standard curve prepared from a ten-fold serial dilution of a quantified and linearized plasmid containing the DNA segment to be amplified.

### Generation of *L. amazonensis* AQP1 overexpressor, knockout and complemented cell lines

To generate *L. amazonensis* parasites expressing AQP1 tagged with EGFP, the gene open reading frame (ORF) was amplified using the primers AQP1-GFP-N-F and AQP1-GFP-N-R for N-terminal fusion (S1 Table). The amplicon was purified and cloned into the *Bam*HI site of the *Leishmania* expression vector pXG-GFP2+ [40]. The construction was transfected by electroporation [41]. Clones were selected in semi-solid M199 containing G418 (20 μg/mL).

For gene deletion, we used the LeishGEdit toolkit based on the CRISPR/Cas9 strategy provided by Prof. Eva Gluenz group [24]. Briefly, we transfected *L. amazonensis* with the plasmid pT007_Cas9_T7 to generate our parental line expressing Cas9 nuclease and T7 RNA polymerase. The online primer design tool www.LeishGEdit.net was used to design primers for amplification of the 5′ and 3′ sgRNA templates and for amplification of donor DNA from pTPuro_v1 plasmid (S1 Table) (Fig. 3a). Following transfection, parasites were plated onto semisolid M199 (1% agar) complemented with puromycin (20 μg/mL) for selection and isolation of knockout clones. For genetic complementation, the *aqp1* ORF fused to EGFP at the N-terminus, was previously cloned into the episomal expression vector pXG-GFP2+ and reintroduced into selected knockout lines by electroporation. Transfectants were selected and cloned in semisolid M199 containing G418 (20 μg/mL).

To confirm replacement of the *aqp1* ORF by the puromycin resistance cassette and *aqp1* episomal complementation, genomic DNA from the selected clones was extracted using the salting out deproteination extraction (SODE) method [42]. The resulting DNA samples were used as template in PCR amplification analyses.

### Cell volume measurements

Cell volume changes in response to hypo-osmotic stress were evaluated as previously described [15, 43], with minor modifications. Briefly, log-phase promastigotes were collected, washed with PBS, and resuspended in a 150 mOsm buffer to a final concentration of 10^9^ promastigotes/mL. Aliquots of 100 μL were transferred to the wells of a microtiter plate. Hypo-osmotic shock was induced by mixing equal volumes of the isotonic cell suspension and deionized water. Absorbance at 550 nm was measured every 15 sec over a 3 min period in a microplate reader (Spectramax 340, Molecular Devices). A decrease in absorbance reflects an increase in cell volume. For isosmotic controls, cell suspensions were diluted with isosmotic buffer without addition of water.

### *In vitro* antimony susceptibility assay

The susceptibility of the *Leishmania* lines of interest to trivalent antimony (Sb^III^) (Sigma-Aldrich) was determined using the MTT [3-(4,5-Dimethylthiazol-2-yl)- 2,5-diphenyltetrazolium bromide] colorimetric assay (Sigma-Aldrich), as previously described [44, 45]. 2×10^6^ log-phase promastigotes were incubated for 24 h at 25°C in 96-well plates with 200 μL of M199 containing increasing concentrations of potassium antimonyl tartrate trihydrate (Sb^III^) (Sigma-Aldrich): 0, 7.81, 15.62, 31.25, 62.5, 125, 250, 500, and 1000 μM. Then, 30 μL of the MTT solution (5 mg/mL) was added and plates were incubated for 3 h at 25°C.

The reaction was stopped by addition of 50 μL of 20% Sodium Dodecyl Sulfate (SDS). Absorbance was measured at 595 nm using a microplate reader (Spectramax 250, Molecular Devices). The 50% effective concentration (EC_50_) was determined using a nonlinear sigmoidal regression model of the concentration-response curves in GraphPad Prism 8.

### Elemental antimony and iron quantification

Inductively coupled plasma mass spectrometry (ICP-MS) was performed as previously described [46]. Briefly, the concentrations of Sb isotope (^121^Sb) and Fe isotope (^56^Fe) were quantified using internal standards and normalized by total protein content of digested promastigote cells. For Sb uptake quantification, 10^8^ *L*. *amazonensis* promastigotes were incubated with 10 μM of potassium antimonyl tartrate trihydrate (Sb^III^) (Sigma-Aldrich) for 15 min and washed three times with PBS. All samples were digested for 1 h in 100 μL of HNO_3_ 70%. Lysates were diluted 10-fold into 1% nitric acid for the analysis.

### Western blot

Approximately 10^7^ promastigotes were lysed with 100 μL of lysis buffer (1% Triton, 150 mM NaCl, 50 mM Tris-HCl pH 7.6) supplemented with a protease-inhibitor cocktail (Roche). Lysates were clarified at 14,000 × *g* for 15 min at 4 °C. Laemmli sample buffer was added to the supernatant and samples were boiled for 5 min. Equal protein amounts (20 μg per lane) were resolved by SDS-PAGE and analyzed by western blot using anti-SODA (1:500) [47], anti-SODB (1:500) [47], anti-FLAG M2 (1:4,000, for Cas9 detection; F1804-MG, Sigma-Aldrich), and anti-arginase (1:500, loading control) [20] antibodies diluted in TBS Blocking Buffer (LI-COR Bioscience). After primary incubation, membranes were washed three times in TBST (TBS, 0.1% Tween-20) and incubated for 1 h at room temperature, protected from light, with IRDye 800CW goat anti-rabbit or anti-mouse (LI-COR Biosciences; 1:6,000 in TBS Blocking Buffer). Blots were washed again in TBST and scanned on an Odyssey CLx imager (LI-COR Biosciences). Band intensities (near-infrared fluorescence) were quantified using NIH ImageJ 1.50i.

### Superoxide dismutase (SOD) activity

Superoxide dismutase (SOD) activity was quantified with the riboflavin–nitroblue tetrazolium (NBT) photoreduction method as described previously [48, 49]. Briefly, 10^8^ promastigotes were washed three times in PBS (KCl 2.7 mM, KH_2_PO_4_ 1.5 mM, NaHPO_4_ 8 mM and NaCl 136.9 mM, pH 7.0) and lysed in 50 mM Tris-HCl pH 7.5 containing 2.5% [v/v] protease inhibitor cocktail (Sigma) by ten cycles of rapid freezing in liquid nitrogen followed by thawing in a 42°C water bath. The lysate was clarified by centrifugation (12,000 × g, 10 min, 4°C) and the protein concentration measured by the Bradford assay [50]. 10 µg of total protein was added to the reaction buffer (50 mM Tris-HCl pH 7.5, 13 mM L-methionine, 75 µM NBT, 0.1 mM EDTA) to a final volume of 200 µL. The reaction was initiated with addition of 4 µM of riboflavin and the mixtures were exposed to white-fluorescent light for 30 minutes at room temperature. Tubes kept in the dark served as blanks, and a riboflavin-containing reaction without lysate represented 100 % NBT reduction. Absorbance was measured at 560 nm, and one unit (U) of SOD activity was defined as the amount of enzyme that inhibits NBT-diformazan formation by 50 % under these conditions.

### *In vitro* mouse macrophage infections

Mouse bone marrow derived macrophages (BMM) were prepared from C57BL/6 mice as previously described [51]. A total of 10^6^ BMMs per well were plated on glass coverslips in 6-well plates and incubated for 24 h at 37 °C 5% CO_2_ in BMM media: RPMI 1640 medium supplemented with 100 U/mL penicillin, 100 μg/mL streptomycin, 5% FBS, and 20% L929 cell-conditioned medium, as a source of macrophage differentiation factors. Infective metacyclic forms were purified from stationary phase promastigote cultures (second to third day after entering stationary phase) using the 3A.1 monoclonal antibody as described [52]. Purified metacyclics were added at a ratio of 5 parasites per macrophage for 4 h at 34 °C. The cells were washed 3 times with PBS and fixed or incubated in BMM media to complete 24, 48, 72 and 96 h of infection. Coverslips were fixed in 4% paraformaldehyde and incubated with 1 μg/ml DAPI for 1 h, after permeabilization with 0.1% Triton X-100 for 10 min. The number of intracellular parasites was determined by counting the total macrophages and the total intracellular parasites per microscopic field (Leica DMI8 inverted fluorescence microscope, Department of Zoology, Institute of Biosciences, University of São Paulo). At least 200 host cells, in triplicate, were analyzed for each time point.

### *In vivo* infection and parasite tissue load determination

Six-week-old female C57BL/6 or BALB/c mice were inoculated as previously described [53] with 10^6^ purified metacyclics from *L. amazonensis* WT, C9/T7, AQP1 KO and add-back (AB) in the left hind footpad in a volume of 0.01 mL. Lesion progression was monitored once a week by measuring the difference in thickness between the left and right hind footpads with a caliper (Mitutoyo Corp., Japan). The parasite load in the infected tissue was determined after 8-12 weeks in the infected tissue collected from footpad lesions of sacrificed mice by limiting dilution [54].

### Phylogenetic Analysis

Putative AQP1 orthologues from trypanosomatid parasites were retrieved from TriTrypDB using the *Leishmania mexicana* AQP1 sequence (LmxM.30.0020) as a query [16]. The database was also searched for putative AQP1 homologues in *Homo sapiens*. Amino acid sequences were aligned in MEGA 12 with MUSCLE (default parameters), and phylogenetic relationships were inferred by Maximum Likelihood (ML) under the Jones–Taylor–Thornton substitution model [55, 56]. The topology with the highest log-likelihood (–5,648.66) was chosen for presentation. Branch support was evaluated by 1,000 bootstrap replicates; percentages of replicate trees in which the associated taxa clustered are indicated at the corresponding nodes [57]. The final data set comprised 13 sequences and 342 unambiguously aligned positions.

### Statistical analysis

Data were analyzed in GraphPad Prism using unpaired, two-tailed Student’s t-tests and one-way ANOVA. Differences were considered statistically significant at p < 0.05. Exact p-values and the number of biological replicates is reported in the corresponding figure legends.

## Acknowledgments

We thank Eva Gluenz (University of Bern) and Stephen Beverley (Washington University) for generous gift of plasmids. We are grateful to Bruno Lemos Batista and Camila Neves Lange (Federal University of ABC) for assistance with ICP-MS analyses; to Mario Costa Cruz for confocal microscopy support at the Core Facility of the Centro de Facilidades para Pesquisa (CEFAP), University of São Paulo; and to Federico David Brown Almeida for providing access to the fluorescence microscope in his laboratory at the University of São Paulo.

## Supporting information captions

**Supplementary Figure 1.** Sequence identity of trypanosomatid aquaglyceroporins. (a) Multiple-sequence alignment of aquaglyceroporins from trypanosomatids. The Pro-Asn-Phe (PNF) peroxisomal-targeting motif is boxed in red. Dots (.) denote residues identical to the reference (top) sequence; dashes (–) indicate gaps. (b) Pairwise percent-identity matrix for the same proteins, calculated as (1 − p-distance) × 100. Protein IDs: *L. amazonensis* LAMAPH8_000653100; *L. mexicana* LmxM.30.0020; *L. donovani* LdBPK_310030; *L. infantum* LINF_310005100; *L. major* LmjF.31.0020; *L. tarentolae* LtaP31.0020; *L. braziliensis* LbrM.00.0079; *Trypanosoma brucei* Tb927.6.1520, Tb927.10.14160, Tb927.10.14170; *Leptomonas pyrrhocoris* LpyrH10_32_1110; *Lept. seymouri* Lsey_0007_0780; *H. sapiens* NP_001161.1.

**Supplementary Figure 2.** Validation of the Cas9/T7-expressing *L. amazonensis* parental line. (a) Western blot for Cas9 in total lysates of wild-type (WT) and Cas9/T7 (C9/T7) *L. amazonensis* promastigotes. L = protein ladder. (b) Growth curves of WT and C9/T7 promastigotes in regular medium (solid lines) vs. heme-depleted medium (dashed lines). Parasite density (cells/mL) is shown as mean ± SEM of 3 independent experiments. (c) MTT viability assays for promastigotes of WT and C9/T7 exposed to increasing Sb^III^ concentrations (0–1000 µM). Percent survival is plotted for each strain. Data are mean ± SEM from 3 independent experiments (each in triplicate). (d) Lesion development in BALB/c mice inoculated with 10^6^ metacyclic promastigotes of WT and C9/T7 lines. Points represent mean footpad thickness (± SEM, n = 5 mice per group) over time. (e) Parasite load in footpad lesions when lesions reached the maximum allowed: 7 weeks post-infection for WT and 9 weeks for C9/T7. Bars show mean ± SEM for 5 mice per group.

**Supplementary Figure 3.** Superoxide dismutase expression in AQP1 mutant lines under heme depletion. (a) Western blots for SOD B (SODB) and SOD A (SODA) in total lysates of WT, KO, and AB promastigotes cultured in control (+) or heme-depleted (-) media. Arginase (ARG) is shown as loading control. (b) Densitometry of SOD bands normalized to ARG. Bar graphs show SODB/ARG and SODA/ARG levels relative to WT. Data represent 3 independent experiments (mean ± SEM).

**S1 Table.** Oligonucleotides used in this study.

